# Unique signals of clinal and seasonal allele frequency change at eQTLs in *Drosophila melanogaster*

**DOI:** 10.1101/2021.07.30.454552

**Authors:** Yang Yu, Alan O. Bergland

## Abstract

Populations of short-lived organisms can respond to spatial and temporal environmental heterogeneity through local adaptation. Local adaptation can be reflected on both phenotypic and genetic levels, and it has been documented in many organisms. Although some complex fitness-related phenotypes have been shown to vary across latitudinal clines and seasons in similar ways in *Drosophila melanogaster* populations, we lack a general understanding of the genetic architecture of local adaptation across space and time. To address this problem, we examined patterns of allele frequency change across latitudinal clines and between seasons at previously reported expression quantitative trait loci (eQTLs). We divided eQTLs into groups by utilizing differential expression profiles of fly populations collected across a latitudinal cline or exposed to different environmental conditions. We also examined clinal and seasonal patterns of allele frequency change at eQTLs grouped by tissues. In general, we find that clinally varying polymorphisms are enriched for eQTLs, and that these eQTLs change in frequency in predictable ways across the cline and in response to starvation tolerance. The enrichment of eQTL among seasonally varying polymorphisms is more subtle, and the direction of allele frequency change at eQTL appears to be somewhat idiosyncratic. Taken together, we suggest that clinal adaptation at eQTLs is distinct than that of seasonal adaptation.

## Introduction

Identifying the evolutionary forces that maintain genetic variation in natural populations remains one of the key questions in population genetics (Gillespie 1998; Casillas and Barbadilla 2017; Charlesworth and Charlesworth 2017). One strong diversifying force is environmental heterogeneity (Dobzhansky 1955; McDonald and Ayala 1974; Gillespie 1998), which can result in balancing selection maintaining genetic variation within a population and species (Levene 1953; Haldane and Jayakar 1963; Charlesworth 2006). Environmental change across the range of many widely distributed species is often associated with latitudinal gradients related to phenology (Viegas et al. 2012; Fjellheim et al. 2014; Kong et al. 2019) and adaptation to temperate environments in general (Bradshaw et al. 2004). These spatially varying aspects of the environment have been shown to drive local adaptation on a genome-wide scale and promote functional diversity (Hancock et al. 2010; Li et al. 2010; Ma et al. 2010; Paaby et al. 2010; Huang et al. 2018; Key et al. 2018). For organisms with short generation times, temporal variation in selection pressures can drive adaptive tracking (Botero et al. 2015). Adaptive tracking has been shown to occur in response to seasonal variation in selection pressures (Dobzhansky and Ayala 1973; Mueller et al. 1985; Rodríguez-Trelles et al. 1996; Ananina et al. 2004; Bergland et al. 2014; Wittmann et al. 2017; Machado et al. 2021), and in principle these adaptive fluctuations across seasons should mirror spatial variation because of common selective pressures imposed by seasonality (Singh and Rhomberg 1987).

Empirical work on *Drosophila melanogaster* has shown similar phenotypic and genetic differentiation in fitness-related traits across a latitudinal cline and between seasons. Lab reared descendants of flies collected in the spring are more starvation tolerant, darker, and show a wider breadth of thermal tolerance, similar to lab reared descendants of flies collected in northern locales (Schmidt et al. 2005, 2008; Schmidt and Paaby 2008). Genetic and genomic work has shown that allele frequency shifts between seasons sometimes shows parallel clinal variation in allele frequency (Bergland et al. 2014; Cogni et al. 2014; Paaby et al. 2014; Machado et al. 2021; Mallard et al. 2018). For instance, candidate adaptive polymorphisms in the gene *couch-potato* show parallel shifts in frequency across space and time: the pro-diapause allele has higher frequency in the spring and in the north, compared to the fall or the south (Cogni et al. 2014); but see (Erickson et al. 2020). Similar patterns are observed for polymorphisms associated with life-history at the Insulin Receptor (Paaby et al. 2014), and polymorphisms associated with immune response (Behrman et al. 2018). However, genome-wide examination of clinal and seasonal changes in allele frequencies reveal that parallel shifts across space and time occur at only ~70% of most strongly differentiated polymorphisms (Machado et al. 2021).

The lack of strong parallelism in allele frequency shifts across space and time in flies could arise from several factors. First, the recent and long-term demographic history of flies collected across a latitudinal cline and between seasons differ (Bergland et al. 2014, 2016). Second, selective forces that vary across latitudinal clines might not exactly mirror seasonal shifts in selection pressure. Finally, the genetic architecture of adaptation across latitude clines might be different from adaptation across seasons. The difference in genetic architecture of seasonal and clinal adaptation could arise from a difference of timescale (Botero et al. 2015; Messer et al. 2016). Clinal adaptation likely occurs over a relatively longer time scale, and across numerous collinear environmental factors (Adrion et al. 2015; Bergland et al. 2016). Seasonal fluctuations in the environment are predictable over the period of months but can be highly unpredictable on shorter timescales. Whether the differences in the timescale of fluctuating selection correspond to differences in the genetic architecture of rapid adaptation remains an open question.

To study the comparative genetic architecture of clinal and seasonal genetic adaptation, we performed an integrative analysis incorporating publicly available gene expression, allele frequency, and eQTL profiles (FlyBase Consortium 1999; Zhou et al. 2012; Bergland et al. 2014; Juneja et al. 2016; Machado et al. 2021; Everett et al. 2020). Gene expression variation has been demonstrated to be important for adaptive evolution in many organisms (King and Wilson 1975; Gompel et al. 2005; Fraser et al. 2010; Richards et al. 2012; Fraser 2013; Mack et al. 2018), allowing us to connect genetic variation with fitness-related traits in a relatively comprehensive framework (Fraser et al. 2011). We partitioned eQTLs into different functional groups based on their association with each of ~4000 genes and novel transcribed regions (NTRs). Genes were divided into functional categories based on signals of local clinal adaptation in gene expression (Juneja et al. 2016), anatomical expression profile (FlyBase Consortium 1999), and plastic changes in expression across a range of environments (Zhou et al. 2012). This functional categorization allowed us to infer aspects of physiological changes underlying adaptation, as many physiological changes are driven by tissue or sex-specific gene expression patterns (Whitehead and Crawford 2005; Bentz et al. 2019).

Our analyses reveal several basic results. First, we show that eQTLs are, in general, enriched among clinally varying polymorphisms but not among polymorphisms that seasonally fluctuate across multiple populations. When we polarize eQTLs based on association with genes that are differentially expressed among latitudinal populations, or in response to several ecologically relevant treatments, clinally varying polymorphisms show predictable shifts in allele frequency. Intriguingly, analysis of seasonal changes in allele frequency within each of 20 paired spring-fall population sets shows that there can be consistent, albeit unpredictable, changes in functionally related eQTLs. Our results support the model that evolution across spatial gradients might be determined by a fundamentally different architecture as evolution across seasons.

## Materials and Methods

### Population allele frequencies and differentiation statistics

We used allele frequency estimates at 1,763,522 SNPs from 46 out of 73 populations as reported by Machado *et al* (2021). This dataset includes paired spring-fall samples from 20 geographically distributed localities across two continents. Machado *et al* (2021) modeled allele frequency change between seasons and across a latitudinal cline along the east coast of North America for each SNP using generalized linear models. We use the output of those models to define “seasonal” and “clinal” polymorphisms based on *p*-value and regression coefficients. We also examined the allele frequency change between spring and fall for each of the 20 population pairs independently.

### eQTL and expression data

We examined allele frequency change across space and time at 72,389 unique eQTLs as identified by Everett *el at* (2020) that are also polymorphic among the 6 clinal and 20 seasonal populations we studied. We grouped them into female-specific, male-specific, and non-sex biased (overlapping) eQTLs. For the 39 out of 159 genes reported as differentially expressed (DE) across the latitudinal cline with known eQTLs (Juneja et al. 2016), we used female-specific (n = 1392) and overlapping (n = 880) eQTL. For the DE genes, with identified eQTLs, that are induced by heat shock (gene = 57), chill coma (gene = 16), starvation (gene = 38), high-temperature (gene = 19) and low-temperature (gene = 20) (Zhou et al. 2012), we used overlapping eQTLs.

### Enrichment analysis of clinal or seasonal SNPs in eQTLs

To calculate enrichment of clinal and seasonal SNPs among eQTLs, we took observed eQTLs and 1000 sets of matched control sites and assessed the number that were clinal or seasonal below a range of *p*-value quantiles for clinality and seasonality. We used these counts to calculate odds-ratios. The control SNPs for eQTLs were matched by chromosome, heterozygosity (binned by 0.05), and shared inversion status of four common inversions (Corbett-Detig and Hartl 2012), as the target eQTLs. We first performed this analysis on all eQTLs, and then performed this analysis using top 5% clinal or seasonal *p*-value quantile as thresholds for the eQTLs associated with each gene.

To gain insight into the physiological basis of local adaptation, we examined the enrichment of eQTLs associated with genes highly expressed in different tissues. We grouped eQTLs by genes that are highly expressed in various tissues based on the FlyBase profiles (FlyBase Consortium 1999), and calculated enrichment by comparing to null distributions generated by control SNPs, as described above. We performed multiple testing using the Bonferroni correction of *p* values *(p* = 0.002) for 23 (for female), 22 (for male) and 20 (for overlap) tests.

### Testing the concordance of allele frequency change across space and time at eQTLs

We tested whether eQTLs change in frequency across space and time in a predictable way (concordance) by combining the sign of allelic effects (i.e., up- or down-regulating) at eQTLs with gene expression data contrasting northern and southern populations (Juneja et al. 2016), or a single population exposed to a variety of ecologically relevant environmental treatments (Zhou et al. 2012).

First, we took genes that are differentially expressed between northern and southern populations and identified eQTLs associated with those genes. For this analysis, we only used female-specific and non-sex-biased (overlapping) eQTLs because Juneja *et al* (2016) measured differential expression only in females. For genes that are more highly expressed in northern populations compared to southern ones, we hypothesized that the eQTL that causes an increase in gene-expression should be more common in northern than southern populations. The converse would be the case for genes that are more highly expressed in southern populations. Fly populations collected in the spring are thought to be genetically hardier than those collected in fall (Bergland et al. 2014), and thus we hypothesized that spring-fall comparisons would mirror north-south comparisons.

We applied a similar approach to genes expressed in response to several environmental treatments (Zhou et al. 2012). For this analysis, we used non-sex-biased eQTLs and genes that do not show sex-treatment interaction based on Zhou *et al* (2012). We focused on differential expression in response to chill-coma, heat-shock, low- or high-temperature exposure, and starvation. We hypothesized that for genes upregulated following chill-coma or low-temperature exposure, the upregulating eQTL alleles associated with those genes will be more common in the north and in the spring, relative to the south or the fall. Conversely, we hypothesized that for genes upregulated following heat-shock and high-temperature exposure, the upregulating allele would be less common in the north and in the spring. We hypothesized that genes upregulated following starvation would follow a similar pattern to chill-coma and low-temperature treatments.

We assigned up- and down-regulating alleles, based on gene expression profiles from Everett *et al* (2020) for each eQTL associated with the identified DE genes from the above two datasets. We incorporated clinal coefficients from a four-population clinal model (CrossPop; Machado *et al* 2021) and calculated allele frequencies for those eQTLs in 3 independent spatial contrasts: Maine vs. Florida (FL-ME), Spain vs. Ukraine (ES-UK), and Pennsylvania vs. Wisconsin (PA-WI). We chose these three pairs of populations because they span the classic latitudinal cline in North America or represent a continental gradient in clinality. For each clinal population, if multi-seasonal samples exist (i.e., spring, fall and frost sample), the allele frequencies were calculated as the averaged allele frequency for multiple seasonal samples.

We also examined seasonal coefficients (CrossPop) and seasonal allele frequency fluctuations in 20 paired spring-fall samples collected in North America and Europe (Machado et al. 2021). We generated a null distribution of expected concordance for each population comparison using matched controls based on chromosome, inversion status, and heterozygosity criteria (binned by 0.05), bootstrapped 1000 times.

We calculated empirical *p*-values for concordance analysis. Let *S* be the observed concordance estimate and *S_0_* be the distribution of concordance estimates from our bootstrap distribution, N is the total number of tests, and defined

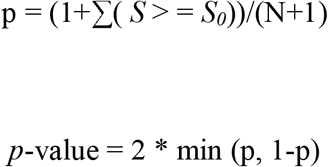

(Davison and Hinkley 1997).

## Results

### Genome-wide Enrichment Test

We find a significant enrichment of female-, male- and non-sex-biased (overlapping) eQTLs among clinal SNPs, but not seasonal SNPs across a range of ranked normalized clinal and seasonal *p*-values (Figure 1A, Supplemental Table 1). After block sampling to reduce to one eQTL per 10kbp, significant enrichment of clinal SNPs in female-, male-and non-sex-biased eQTLs still hold across certain ranges of clinal *p*-value thresholds (Figure S1).

**Figure 1.**
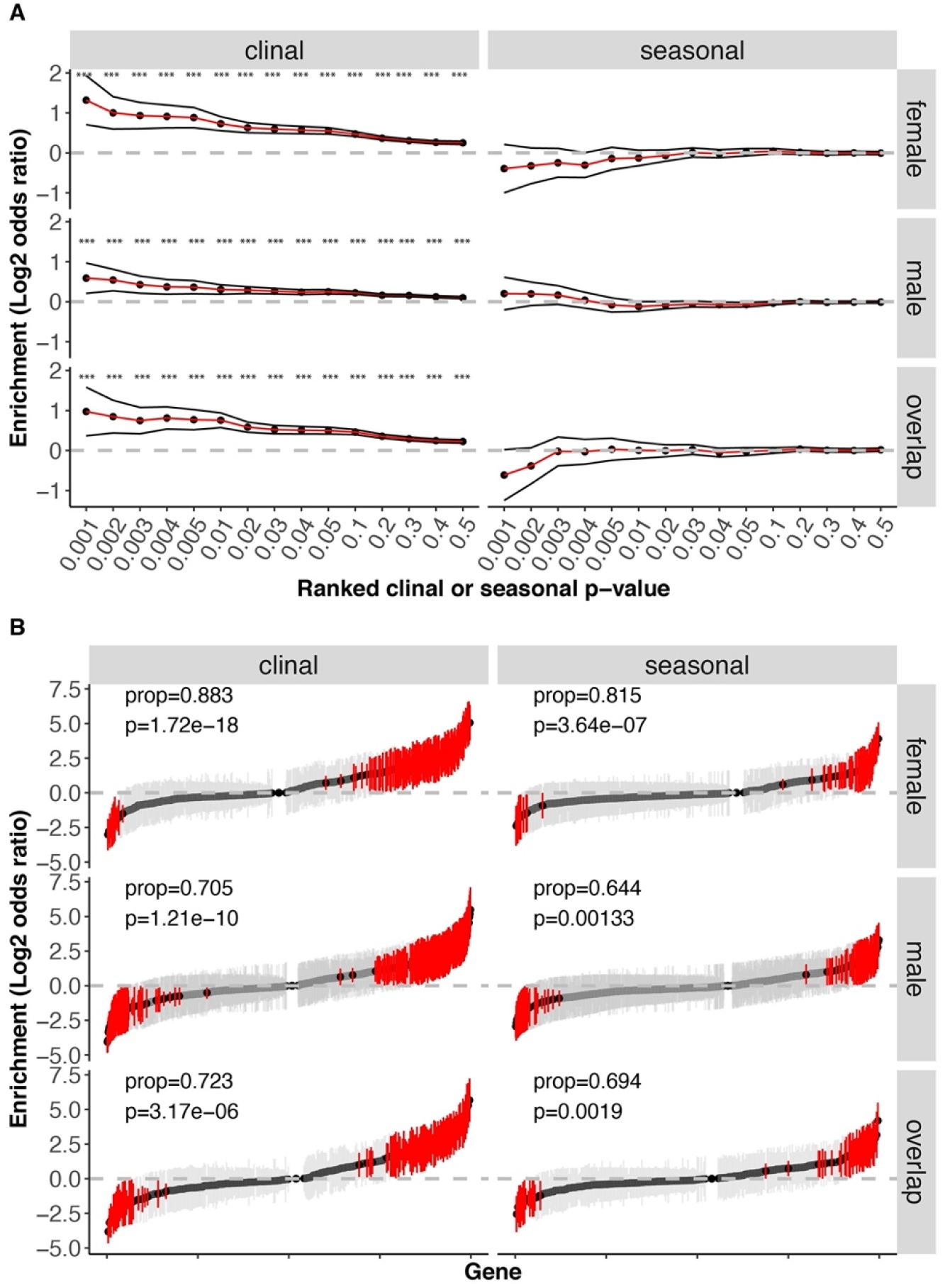
Enrichment of clinal or seasonal SNPs in female-specific, male-specific and overlapping eQTL genome-wide (A), and in every gene identified with eQTL (B). (A) The x-axis is ranked clinal (left) or seasonal (right) *p* value thresholds. The y-axis is enrichment, calculated as the log2 odds ratio of eQTLs having ranked clinal or seasonal *p* values below or equal to certain thresholds compared to matched controls based on matching parameters. Black dots represent average log2 odds ratio over 1000 bootstraps. Black lines are confidence intervals, represented by 1.96 standard deviations of the mean over 1000 bootstraps. Asterisks indicate significant enrichment (*p* = 0.05). (B) The x-axis is genes identified with eQTLs. Genes are ranked by averaged log2 odds ratio within each analysis type (clinal or seasonal) and sex (female, male, overlap) combination panel. The y-axis is enrichment. Black dots represent average log2 odds ratio over 1000 bootstraps. Red or grey error bars are 1.96 standard deviations of the mean over 1000 bootstraps for genes with significant or insignificant signals, respectively. Proportion (prop) represents the ratio between genes significantly enriched for clinal or seasonal eQTLs and the total number of genes with significant (enrichment and depletion) signals.

Next, we tested whether specific genes in our dataset show strong signals of enrichment for clinal or seasonal SNP by calculating an enrichment score for eQTLs associated with each gene relative to unlinked matched controls. We find that many genes (25.879%) are significantly enriched for clinal eQTLs (Figure 1B). Moreover, there is an excess of genes which are enriched for clinal eQTLs, compared to genes which are depleted for clinal eQTLs in females (proportion = 0.883, *p* = 1.72e-18), males (proportion = 0.705, *p* = 1.21e-10), and overlapping (proportion = 0.723, *p* = 3.17e-06) genes and novel transcribed regions (NTRs). Similarly, there is an excess of genes which are enriched for seasonal eQTLs, compared to genes which are depleted for seasonal eQTLs in females (proportion = 0.815, *p* = 3.64e-07), males (proportion = 0.644, *p* = 0.00133), overlapping (proportion = 0.694, *p* = 0.0019) genes and NTRs for female-specific, male-specific and overlapping eQTLs, respectively (Figure 1B).

### Enrichment signal of clinal and seasonal eQTLs in tissues

We asked if enrichment signals of clinal or seasonal SNPs differ based on the reported expression level of genes in different tissue types. We identified eQTLs that are associated with genes that have “high” expression level in each tissue (FlyBase Consortium 1999). We show that clinal eQTLs tend to be associated with genes highly expressed in tissues such as adult head and larval trachea (non-sex-biased eQTL) and are significantly depleted among genes highly expressed in larval hindgut or midgut (Figure 2, Supplemental Table 2). In general, we observe patterns of depletion of seasonal eQTLs across all tissue types (Figure 2).

**Figure 2.**
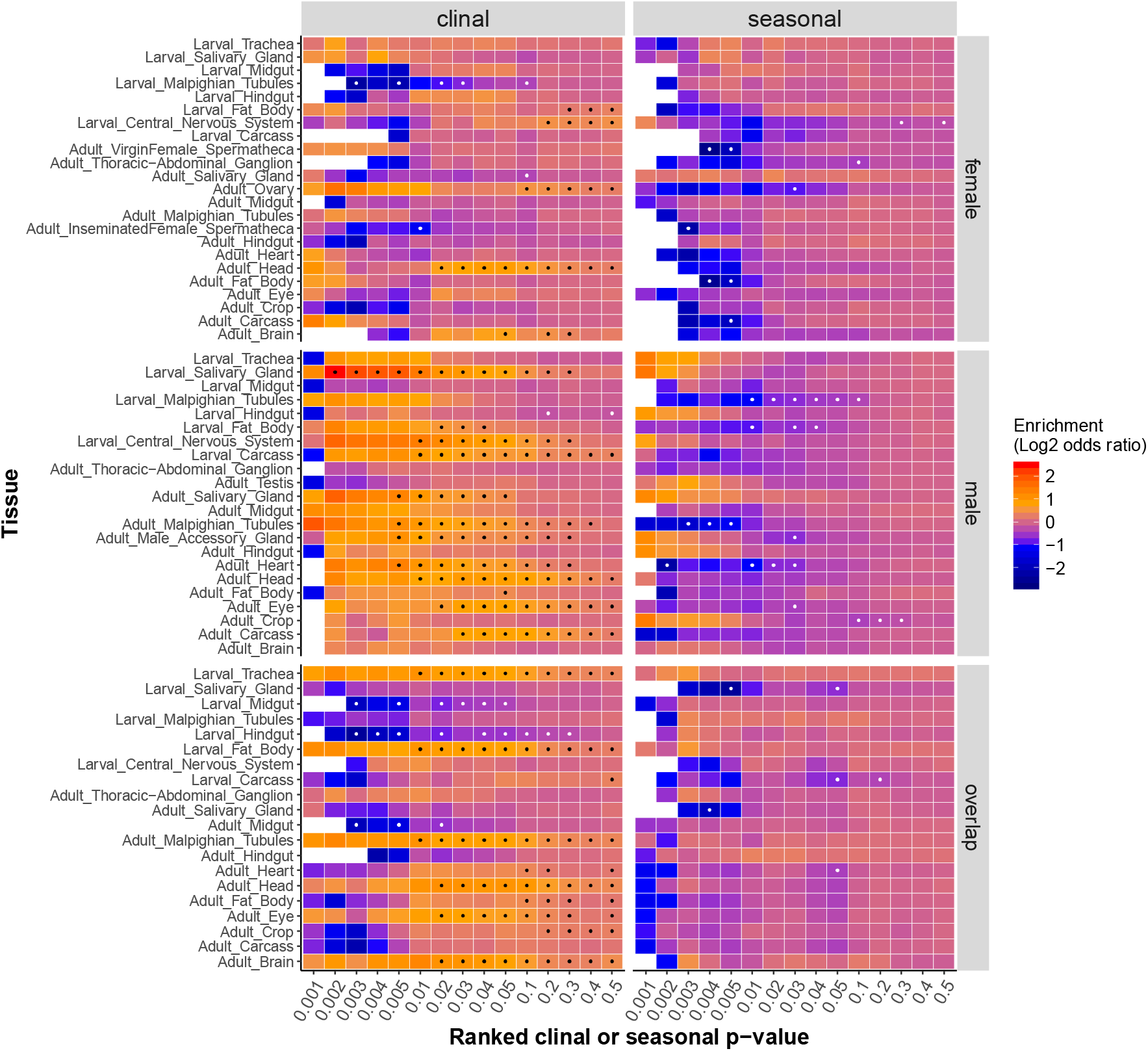
Enrichment of clinal or seasonal SNPs in female-specific, male-specific and overlapping eQTLs, grouped by high expression genes in tissues. The x-axis is the ranked clinal (left) or seasonal (right) *p* value thresholds. The y-axis is tissues. Enrichment is calculated as the log2 odds ratio of eQTLs, grouped by high expression genes in certain tissues, having ranked clinal or seasonal *p* values below or equal to certain thresholds compared to matched controls. Black or white dots indicate significance for enrichment or depletion over 1000 bootstraps, with Bonferroni correction of empirical *p* values (*p* = 0.002) for 23 (for female tissues), 22 (for male tissues), or 20 (for overlapping tissues) tests.

### The concordance of eQTL frequency change across space and time

We tested whether eQTLs show predictable changes in allele frequency across the cline and between seasons. We show that female-specific eQTLs associated with latitudinal DE genes are more likely to change allele frequencies in the predictable direction than controls are (CrossPop: p<0.001; FL-ME: p<0.001; ES-UK: p<0.001; PA-WI: p<0.001; Figure 3A). The same general pattern is observed in overlapping eQTLs across the North American latitudinal cline (CrossPop) and for Florida-Maine (FL-ME) comparisons, but not for Spain-Ukraine (ES-UK) or Winsconsin-Pennsylvannia (PA-WI) comparison. We also show opposite concordance signals for female-specific eQTL associated with latitudinal DE genes in seasonal comparison (CrossPop: p<0.002), but not for overlapping eQTL (Figure 3A). In addition, the inconsistent signals amongst 20 spring and fall comparisons suggest that the eQTLs associated with latitudinal DE genes are not always changing allele frequencies in predicated directions in every seasonal sample (Figure 3A).

**Figure 3.**
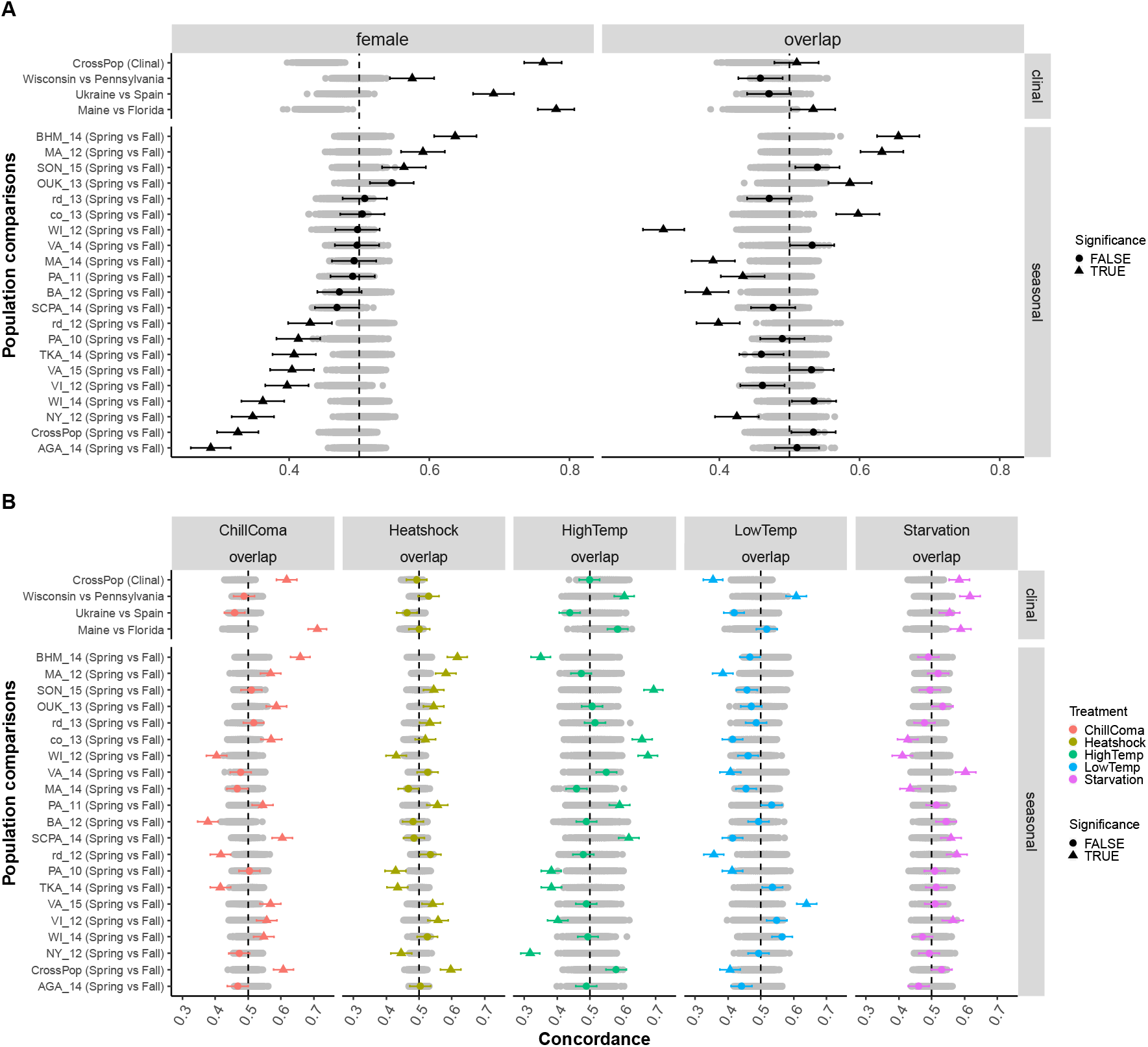
The concordance of allele frequency change for female-specific and overlapping eQTLs associated with genes differentially expressed between high and low latitudinal populations (A), or for overlapping eQTLs associated with genes differentially expressed under certain environmental treatments (B). The x-axis is the fraction of concordance SNPs. The y-axis is clinal or seasonal population comparisons, with the assumption that gene expression patterns are similar between northern and spring populations, and similar between southern and fall populations. The cross-population comparison results were generated by using clinal or seasonal coefficients while other comparisons used allele frequencies of eQTLs from each sample pair. Black (A) or colored (B) dots (triangles and circles) are observed values of eQTLs. Grey circles are expected distributions generated by matched control SNPs, bootstrapped 1000 times. Black (A) or colored (B) triangles indicate observed fractions of concordance eQTLs significantly deviate from expected matched control distributions, with Bonferroni correction of empirical *p* values for 25 tests (*p* = 0.002). Black (A) or colored (B) circles indicate non-significant deviations of observed eQTLs values from expected matched control distributions.

Next, we evaluated the concordance of eQTL allele frequency change across space and time at genes which are differentially expressed between environmental treatments. Consistent with our prediction that northern flies are more starvation tolerant and more chill coma resistant, eQTLs associated with DE genes under starvation (CrossPop: p<0.001, FL-ME: p<0.001, ES-UK: p<0.002; PA-WI: p<0.001; Figure 3B) and chill coma (CrossPop: p<0.001, FL-ME: p<0.001; Figure 3B) treatments show concordance signals in clinal comparisons. In contrast to our prediction that northern flies are more low temperature adapted, eQTLs associated with low temperature treatment induced DE genes show opposite concordance signal for clinal comparison (CrossPop: p<0.002, Figure 3B). We show concordance signals for eQTLs associated with chill coma (CrossPop: p<0.001, Figure 3B) and heat shock (CrossPop: p<0.001, Figure 3B) treatment induced DE genes in seasonal comparisons, consistent with our prediction that spring flies are more chill coma and heat shock resistant, but opposite concordance signal for eQTLs affecting low temperature induced DE genes (CrossPop: p<0.002, Figure 3B). Similar to results for eQTLs associated with latitudinal DE genes, concordance signals for eQTLs under environmental treatments are inconsistent amongst 20 seasonal spring and fall comparisons (Figure 3B).

We also evaluated whether a limited number of genes, with more eQTLs identified than others, are driving the concordance signal. We find that the significant clinal SNP concordance signal of eQTLs affecting known latitudinal DE genes is mainly driven by two genes, *AANATL3* and *Hsc70-2*. In particular, *Hsc70-2*, a heated shock related gene harboring ~1000 eQTLs, changes allele frequencies in opposite directions between clinal and seasonal comparisons under the assumption that spring population is similar to northern population (Figure S2).

### The concordance of allele frequency change for two genes

To further study the clinal/seasonal eQTL allele frequency patterns of two fitness-related genes, we evaluated whether the allele frequency of *Ace* changes in parallel with *Hsc70-2* in both clinal and seasonal comparisons. We show that although within the same region of recent soft sweep (Garud et al. 2015), the allele frequency concordance signal of the two genes is in parallel in seasonal comparison but not in clinal comparison. This suggests that selection can act independently on these two genes, resulting in different allele frequency change patterns.

## Discussion

Although there are well documented patterns of local adaptation across latitudinal clines and between seasons in *D. melanogaster*, we still lack a comprehensive genome-wide understanding of the genetic architecture of this adaptive evolutionary change. Here, we performed an analysis combining eQTLs, gene expression profiles, and patterns of genetic differentiation among populations to infer aspects of the genetic architecture of local adaptation. We define genetic architecture broadly to encompass features of physiology, anatomy and sex. Our results show that the genetic architecture of clinal and seasonal adaptation differ at eQTLs, suggesting that distinct evolutionary processes do occur across space and time in this species.

### Enrichment

We show that eQTLs are, in general, enriched for clinally varying polymorphisms, but less so for seasonally varying ones (Figure 1, Supplemental Table 1). The levels of enrichment of clinal polymorphisms observed here are larger than those observed for other functional categories in *Drosophila* (Machado et al. 2016) and other species (Ye et al. 2013; Mack et al. 2018), suggesting that SNPs underlying clinal adaptation are more likely to be eQTLs than other categories. This finding agrees with growing literature showing evidence that spatial differentiation at eQTLs contributes to local adaptation across various taxa (Fraser 2013; Gould et al. 2017; Kitano et al. 2018; Mack et al. 2018; Phifer-Rixey et al. 2018; Colicchio et al. 2020).

By grouping eQTLs based on tissue type, we can gain insight into the physiological basis of local adaptation (Larsen et al. 2013; Porcelli et al. 2016). We observe that clinally varying polymorphisms are significantly enriched among eQTL associated with genes highly expressed in trachea, the fat body, spermatheca, ovary, and head (Figure 2). These tissues are important for respiration (Hayashi and Kondo 2018), nutrient storage (Zheng et al. 2016), and reproduction (Nonidez 1920; Zhao et al. 2016; Erickson et al. 2020) and could plausibly underly life-history trade-offs observed among populations sampled across the latitudinal gradients (Schmidt et al. 2005; Klepsatel et al. 2013). Intriguingly, we find a significant deficit of clinal eQTLs associated with genes highly expressed in the larval gut system, suggesting that these tissues could be under stronger evolutionary constraint in the early developmental stages (Salvador-Martínez et al. 2017).

We hypothesized seasonally varying polymorphisms would be enriched for eQTLs in similar ways as clinally varying polymorphisms are if seasonal and clinal adaptation had similar genetic architectures (Rhomberg and Singh 1988; Machado et al. 2021; Erickson et al. 2020; Rodrigues et al. 2020). However, we do not observe a strong signal of enrichment of seasonal polymorphisms at eQTLs, like we do for clinal polymorphisms (Figures 1–2, Supplemental Tables 1-2). First, we show that there is no enrichment of seasonal SNPs in eQTLs, genome-wide (Figures 1, S1), highlighting the challenge of functional prediction of previously reported seasonal SNPs in *D. melanogaster* populations (Bergland et al. 2014; Machado et al. 2021). Second, we show that there is a general depletion of seasonal SNPs in eQTLs associated with various tissue types. The depletion patterns indicate that there are idiosyncratic changes between populations for spring-fall samples (see *Concordance of eQTLs*, below).

We observe a number of tissues showing differences in enrichment signals between clinal and seasonal eQTLs. For example, we show significant enrichment of clinal SNPs in eQTLs affecting high expression genes in adult head (Figure 2). This is consistent with a previous study showing that DE genes in the heads of flies that show diapause, a life-history trait known to be important for local adaptation, are clinal (Zhao et al. 2016). However, those same eQTLs are not enriched for seasonal SNPs, inconsistent with previous expectations (Zhao et al. 2016). The discrepancy between our result and previous findings could be affected by the genes that are highly expressed in multiple tissues (Müller et al. 2011), which may affect multiple physiological traits under selection. Different from results shown by Zhao et al (2016), we observe a significant enrichment signal of clinal eQTLs in the female ovary. This is in line with the widely accepted understanding that reproductive tissues represent important physiological basis for local adaptation (Ellegren and Parsch 2007). Additionally, when grouping overlapping eQTLs by high expression genes in larval trachea, which is involved in respiration adaptation (Hayashi and Kondo 2018), we show that they are enriched for clinal SNPs but not seasonal ones (Figure 2). This result does not agree with a previous study on candidate genes showing that major effect genes affecting respiration adaptation harbor both clinal and seasonal SNPs (Mallard et al. 2018). Many of those studies focus on identifying a limited number of shared genetic loci underlying both clinal and seasonal genetic adaptation, but they may be biased from a genome-wide perspective (Cogni et al. 2014; Mallard et al. 2018).

### Concordance of eQTLs

We examined whether there are changes in allele frequency across space and time that are predictable based on the sign of allelic effect and phenotypic differentiation amongst populations sampled along the cline or through the growing season. Concordant signals of allele frequency change at eQTLs between clinal or seasonal populations is indicative of genetic adaptation underlying adaptive DE genes (Bergland et al. 2014; Machado et al. 2021; Rodrigues et al. 2020). For instance, for genes that have higher expression levels in northern population than in southern population, we expect the eQTLs that up-regulate those genes to be more common in northern population. We show that eQTLs grouped by previously reported latitudinal DE genes change allele frequencies concordantly (in predictable directions) across a latitudinal cline in North America and in Europe (Figure 3). Such result suggests that latitudinal gene expression changes are driven by allele frequency changes of regulatory regions, indicating genetic adaptation. Similar clinal concordance signal is observed for eQTLs associated with DE genes induced by starvation treatment (Figure 3), which agrees with previous findings on northern flies to be more stress-tolerant than southern flies (Schmidt et al. 2005; Arthur et al. 2008). The clinal concordance of eQTLs from our study is in line with recent work showing direct evidence of predictable eQTL allele frequency change underlying spatial adaptive evolution in several species including fish (O’Brown et al. 2015; Jacobs et al. 2020), coral (Rose et al. 2018), and human (Fraser 2013). By incorporating various treatment induced DE genes, this finding has also broadened our view in studying local adaptation of *Drosophila* focusing a small number of functional loci (Cogni et al. 2014; Paaby et al. 2014; Behrman et al. 2018) to a more general scope.

We observe changes in allele frequency between seasonal eQTLs that are idiosyncratic. For example, when grouping eQTLs by latitudinal DE genes, only three (female) and four (overlapping) out of twenty seasonal population comparisons show strong concordance signals at eQTLs, respectively. Eight (female) and six (overlapping) comparisons show strong opposite signals, suggesting that in these seasonal populations, eQTLs are more likely to change allele frequencies in opposite directions than expected (Figure 3A). These results suggest a distinct genetic basis of evolutionary change across space and time. We observe similar results of idiosyncratic seasonal concordance signals when grouping eQTLs by treatment-specific DE genes (Figure 3B). The idiosyncractic seasonal change in allele frequency among clines and seasons is similar to patterns observed by Erickson *et al* (2020) that examined diapause associated SNPs. Idsioyncratic differences in the direction of allele frequency change could be due to subtle differences in weather (Machado et al. 2021), polygenic adaptation (Barghi et al. 2019), or a complicated short-term demographic event.

We show that the eQTL allele frequency patterns of two neighboring genes, *Ace* and *Hsc70-2*, are different between clinal and seasonal adaptation (Figure 4). These two genes are physically close (open reading frame distance: ~0.176Mbp) and located in an region that has been identified under recent soft sweep on chromosome 3R (Garud et al. 2015). We show that the predicated change of expression levels for the two genes are similar from spring to fall, but different from north to south. Such result suggests that the two fitness-related genes are independently selected, and selection forces could be different between clinal and seasonal adaptation.

**Figure 4.**
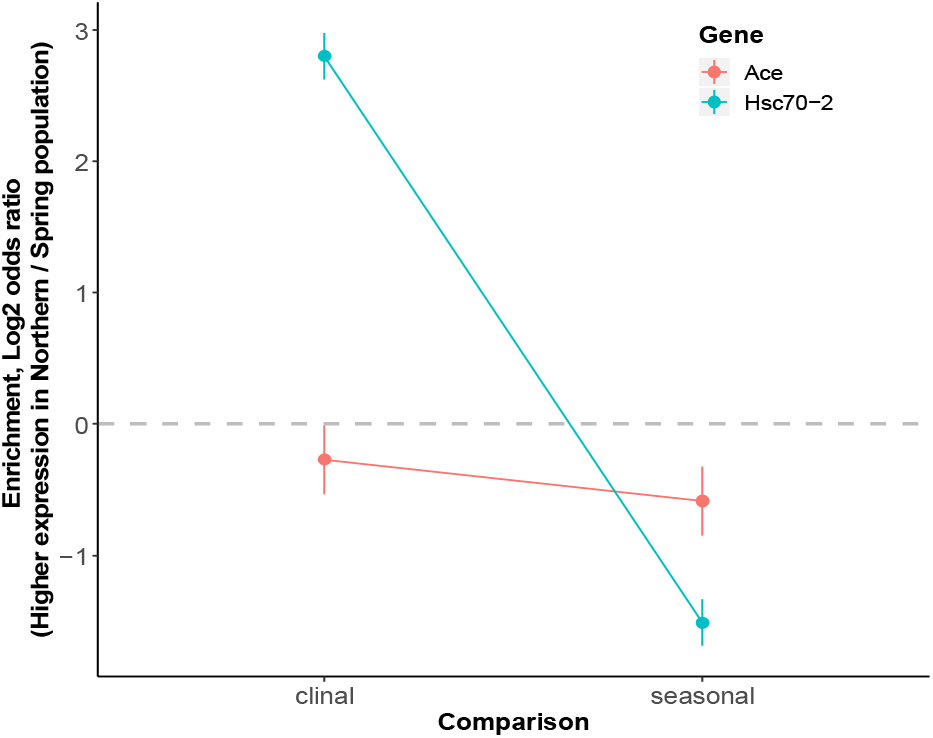
Predicted expression of *Ace* and *Hsc70-2* based on concordance of eQTL allele frequency. The x-axis is clinal or seasonal comparisons using cross-population model clinal or seasonal coefficient values, respectively. The y-axis is enrichment, calculated as the log2 odds ratio of eQTL changing allele frequency in the expected direction compared to matched controls based on matching parameters. An enrichment value greater than 0 indicates higher expression levels in northern or in spring populations whereas an enrichment value smaller than 0 indicates higher expression of genes in southern or in fall populations. Dots represent average log2 odds ratio over 1000 bootstraps. Error bars are confidence intervals, represented by 1.96 standard deviations of the mean over 1000 bootstraps.

### Conclusions

Taken together, we suggest that seasonal adaptation at eQTLs may be idiosyncratic amongst populations. The idiosyncratic patterns in our tests for seasonal eQTLs (Figure 3) suggest that seasonal adaptation cannot be explained solely by a common set of seasonally oscillating SNPs. We propose two possibilities explaining the idiosyncratic patterns for seasonal adaptation. First, the architecture of genetic variation in ecologically relevant traits could be under diverse selection pressures in different geographical population-specific seasonal environments (Erickson et al. 2020). Therefore, eQTLs affecting a trait that has overall fitness advantage in one spring-fall seasonal comparison in one geographic location might not be favored in another, resulting in inconsistent allele frequency changes amongst populations. This possibility is supported by our results on seasonal population-specific concordance signals (Figure 3). Second, seasonal adaptation could be achieved by a subset of common eQTLs via combinations with other population-specific seasonal loci amongst populations. It has been shown in previous studies that different combinations of genetic loci in replicate populations could evolve to adapt to the same selective condition while only a limited number of common loci is identified (Barghi et al. 2019). Such possibility could explain the lack of genome-wide enrichment signals of seasonal SNPs in eQTLs (Figures 1, S1), and in high-expression genes in tissues (Figure 2). To better understand the genetic architecture of local adaptation from a gene expression perspective, more clinal and seasonal DE genes and eQTLs need to be identified in future works.

## List of supplemental figures and tables

Supplemental Figure S1: Enrichment of clinal or seasonal SNPs in female-specific, and male-specific and overlapping eQTLs after block sampling.

Supplemental Figure S2: *AANATL3* and *Hsc70-2* drive concordance signal of eQTLs grouped by latitudinal DE genes.

Supplemental Table 1: Summary of genome-wide enrichment analysis for eQTLs.

Supplemental Table 2: Summary of enrichment analysis for eQTLs grouped by tissues.

**Figure S1.**
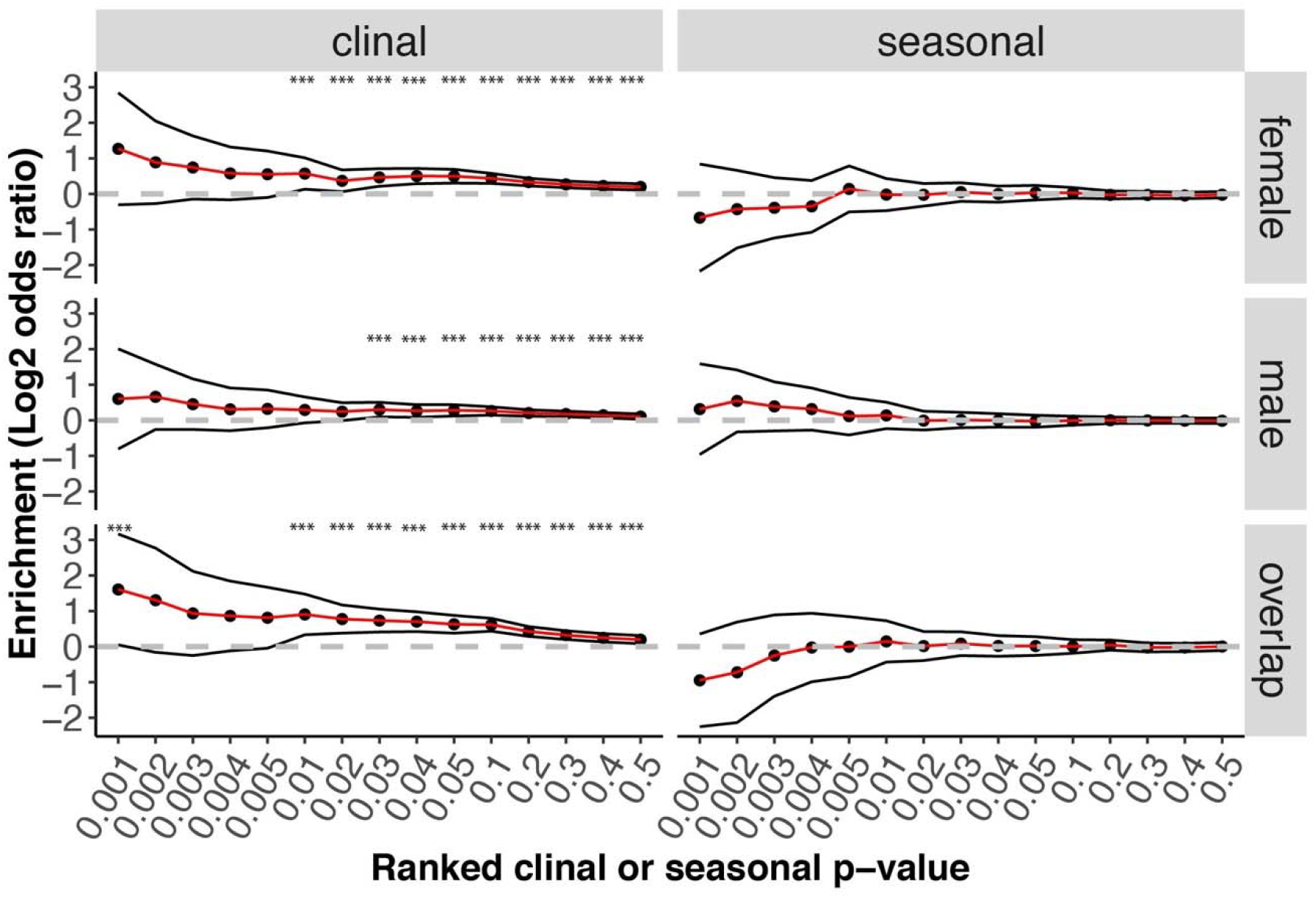
Enrichment of clinal or seasonal SNPs in female-specific, and male-specific and overlapping eQTLs after block sampling. The x-axis is ranked clinal (left) or seasonal (right) *p* value thresholds. The y-axis is enrichment, calculated as the log2 odds ratio of eQTLs having ranked clinal or seasonal *p* values below or equal to certain thresholds compared to matched controls. Female-specific, or male-specific and overlapping eQTLs are down-sampled 100 times with only 1 random eQTL selected in every 10kb non-overlapping windows for each downsampling. Black dots represent average log2 odds ratio across 100 samples over 1000 bootstraps each. Black lines are confidence intervals (1.96 standard deviation of the mean). Asterisks indicate significant enrichment *(p* = 0.05).

**Figure S2.**
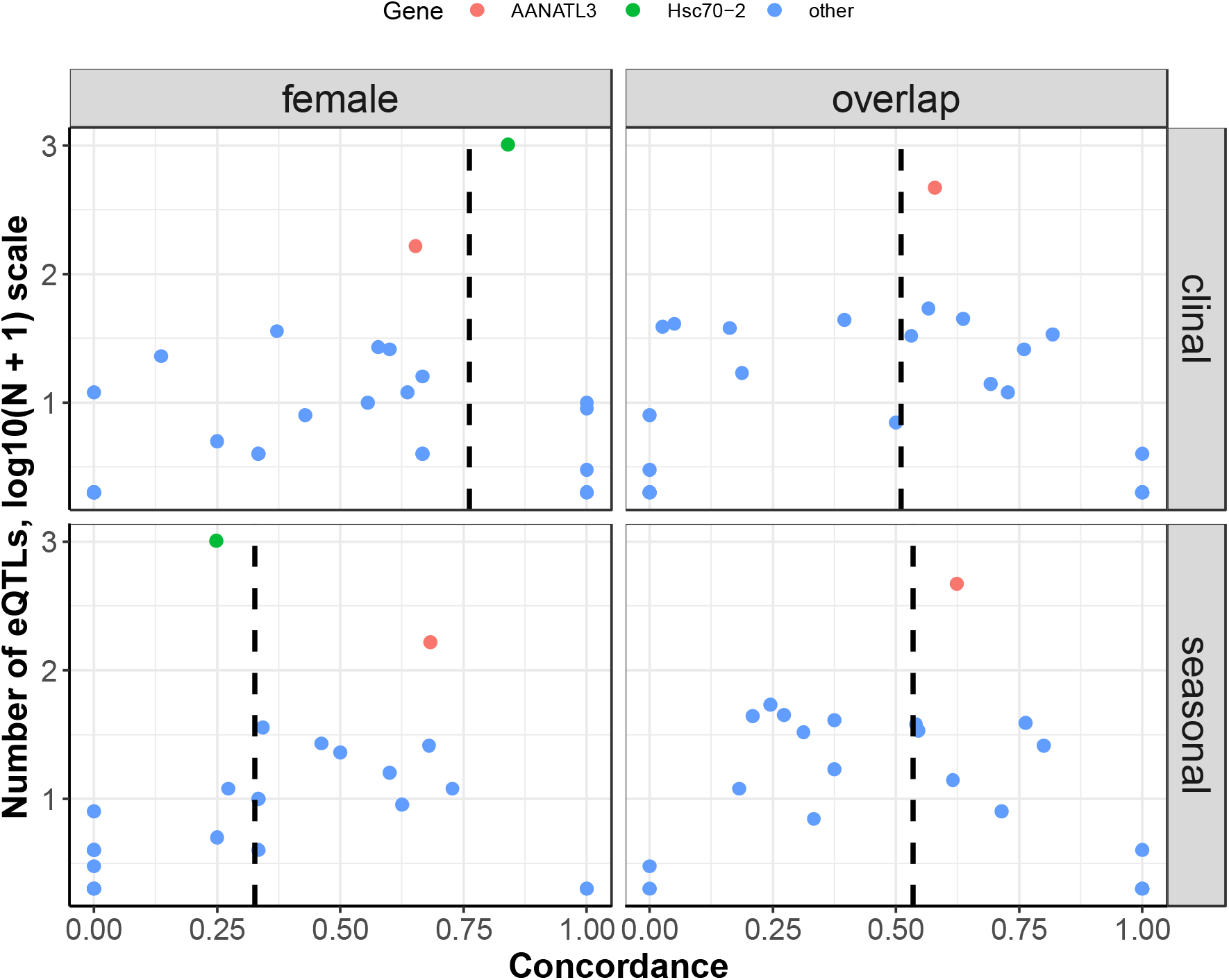
*AANATL3* and *Hsc70-2* drive concordance signal of eQTLs grouped by latitudinal DE genes. The x-axis is the fraction of concordance eQTLs grouped by latitudinal DE genes. The y-axis is the number of eQTLs (log10(N+1) scale) identified with each gene. The black dashed lines are the cross populational (CrossPop) values.

